# A neuroendocrine principle: Pancreatic islets actively shape sympathetic innervation

**DOI:** 10.1101/2025.09.03.673215

**Authors:** Dong-Xue Li, Jia-Mei Luo, Jun-Jie Wang, Yun-Zhen Qian, Dilinazi Abudujilile, Musitaba Mutailifu, Tian Yang, Yuan-Xin Hong, Wen-Tao Shi, Xiao-Yi Ma, Qing Ye, Lei Zhu, Hui-Li, Xiao-Mei Yang, Yan-Li Zhang, Shu-Heng Jiang, Yan-Qiu Yu, Kai Wang, Jun Li, Qing Li, Li-Peng Hu, Xue-Li Zhang, Zhi-Gang Zhang

## Abstract

Survival critically depends on maintaining blood glucose levels to provide essential energy, especially during emergencies such as the fight-or-flight response, when timely glucose control via neural integration is vital. However, pancreatic islets constitute only a small fraction of the pancreas and are dispersed throughout the organ, raising the fundamental question of how the nervous system coordinates synchronized control of multiple islets. Using whole-organ clearing and 3D imaging, we mapped pancreatic sympathetic innervation, revealing specialized anatomical integration between sympathetic nerves and islets. Transplanted islets intrinsically attracted sympathetic nerves independent of their native environment. Chronic islet injury disrupted sympathetic innervation and markedly impaired nerve regeneration after denervation. Sympathetic denervation markedly elevated islet-derived Reg2 and Reg3β; administration of these proteins accelerated sympathetic regeneration and improved islet graft function. Our findings identify an islet-sympathetic architecture actively maintained by islets, uncovering an endocrine-driven mechanism for neural regulation, highlighting Reg2 and Reg3β as therapeutic candidates for diabetes management.

## Introduction

Precise coordination between the nervous and endocrine systems is essential for maintaining metabolic homeostasis and enabling rapid physiological adaptation^1–6^. Pancreatic islets, highly specialized endocrine micro-organs that tightly regulate blood glucose levels, depend heavily on neural inputs to rapidly and dynamically adjust hormone secretion in response to fluctuating metabolic demands^7–14^. In particular, during acute metabolic emergencies such as hypoglycemia or the fight-or-flight response, timely and synchronized regulation of pancreatic islets by the sympathetic nervous system is crucial for restoring blood glucose levels and thus ensuring survival. However, given their dispersed distribution within the pancreas, how sympathetic nerves coordinate rapid and synchronized control of multiple islets under these critical conditions remains unclear. This suggests a specialized neural-islet architecture supporting such coordinated control — an anatomical relationship conventional two-dimensional histological methods cannot resolve at the whole-organ scale^15–20^.

Recently, advances in whole-organ tissue clearing combined with 3D imaging have opened new possibilities to address these critical technical challenges, enabling investigations into islet microstructure, vascular networks, and neural innervation at unprecedented spatial resolution^21–27^. Nevertheless, due to the inherently soft, anatomically complex, and highly convoluted nature of mouse pancreatic tissue, precisely mapping the global 3D structural relationships between islets and neural networks remains technically challenging. Here, we overcame these technical limitations by fully unfolding the pancreas and employing whole-organ tissue clearing combined with advanced 3D imaging. This enabled us to identify an intimate structural association specifically between sympathetic nerves and pancreatic islets, a relationship notably absent for parasympathetic and sensory nerves.

Building on this newly identified anatomical relationship, we further explored its underlying mechanisms. Developmental studies have previously demonstrated that disruption of sympathetic innervation during early pancreatic development severely impairs islet architecture and profoundly compromises hormone secretion^28,29^, reinforcing the classical view of islets as passive recipients of neural inputs whose structural and functional integrity critically depends on innervation. Extending this classical view, our findings provide evidence that the intimate structural relationship between sympathetic nerves and pancreatic islets is actively established and maintained by the islets themselves. These results significantly expand our understanding of endocrine-neural interactions, highlighting an active role for islets in shaping their neural environment.

## Results

### 1. Pancreatic islets are intimately and regularly associated with sympathetic nerves

We first performed a robust method to fully unfold the convoluted mouse pancreas (Extended Data Fig.1a), enabling efficient whole-pancreas tissue clearing (Extended Data Fig.1b). Subsequent advanced 3D imaging provided unprecedented visualization of pancreatic neural networks and their precise spatial relationships with pancreatic islets. To comprehensively characterize the innervation pattern, we specifically labeled sympathetic nerves with tyrosine hydroxylase (TH), parasympathetic nerves with vesicular acetylcholine transporter (VAChT), and sensory nerves with calcitonin gene-related peptide (CGRP). These nerves were labeled in magenta fluorescence, while islets were labeled with insulin (green) and nuclei with DAPI (blue).

Strikingly, sympathetic nerves exhibited a highly organized architecture analogous to “electrical wiring”, characterized by a prominent central conduit traversing the pancreas and numerous finer branches extensively distributed throughout the tissue (Fig. 1a; Supplementary Video 1). Pancreatic islets were not randomly scattered but precisely positioned along these sympathetic nerve fibers, reminiscent of “light bulbs” systematically arranged along “electrical cords” (Fig. 1a-d). High-resolution imaging confirmed intimate spatial contacts between sympathetic fibers and pancreatic islets (Fig. 1e), clearly illustrated by 3D reconstructions (Fig. 1f, g). This anatomical organization might function similarly to “electrical wiring” powering synchronized illumination of connected “bulbs”, enabling islets to simultaneously respond to sympathetic signals. We also examined parasympathetic and sensory innervation in relation to pancreatic islets but did not observe the structured “wiring-light bulb”-like arrangement seen with sympathetic nerves (Extended Data Fig.2; Supplementary Videos 2, 3).

**Figure 1.**
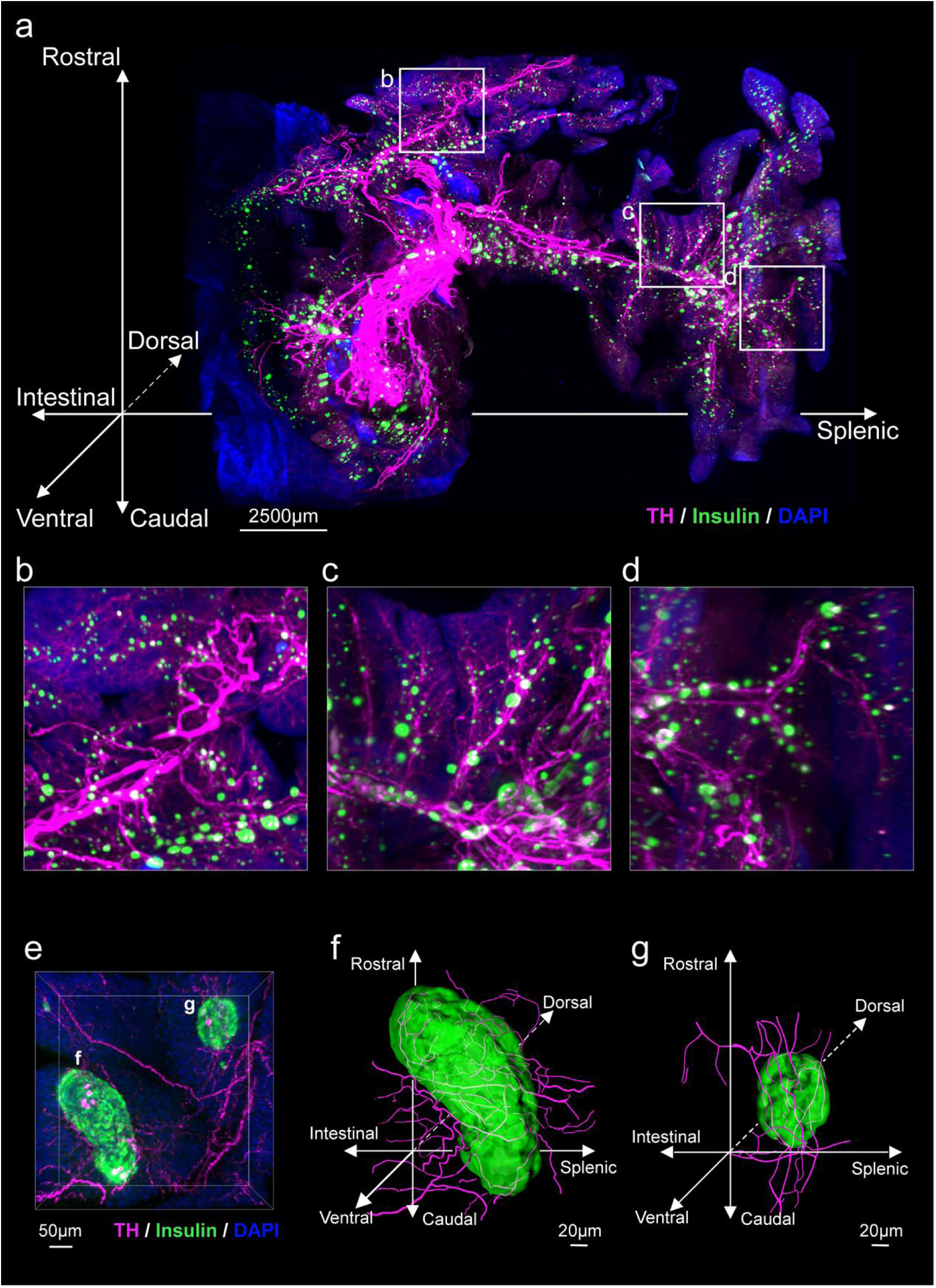
Pancreatic islets form a highly organized spatial arrangement along sympathetic nerves. **a**, Whole-pancreas 3D imaging of an unfolded and cleared pancreas from an 8-week-old male mouse (C57BL/6J) reveals a highly organized sympathetic neural network (TH, magenta). Pancreatic islets (insulin, green) are precisely positioned along these sympathetic nerve branches. Nuclei are stained with DAPI (blue). Scale bars: 2500 μm. See Supplementary Video 1. **b-d**, Magnified views of representative areas from (a). **e,** High-resolution imaging of a pancreatic islet and sympathetic nerve fibers. Scale bars: 50 μm. **f, g,** 3D reconstructions of the two individual islets shown in (e). Scale bars: 20 μm.

The unique anatomical arrangement between islets and sympathetic nerves likely supports their coordinated biological functions. However, it remains unclear how this pattern arises — whether determined by the pancreatic environment or actively shaped by islets themselves.

### 2. Islets recruit sympathetic nerves independently of the pancreatic environment

To distinguish between these two possibilities, we isolated approximately 500 islets from healthy 8-week-old mice and transplanted them beneath the renal capsule of streptozotocin (STZ)-induced diabetic mice^30,31^, establishing a heterotopic transplantation model (Fig. 2a). Using this system, we rigorously tested whether islets could intrinsically establish similar spatial relationships with sympathetic nerves independent of their native pancreatic environment. Post-transplantation monitoring confirmed graft viability and restored endocrine function, indicated by normalized blood glucose levels (Extended Data Fig. 3b, e). Notably, the model employed here was primarily intended to investigate biological mechanisms rather than to serve as a therapeutic approach for diabetes.

**Figure 2.**
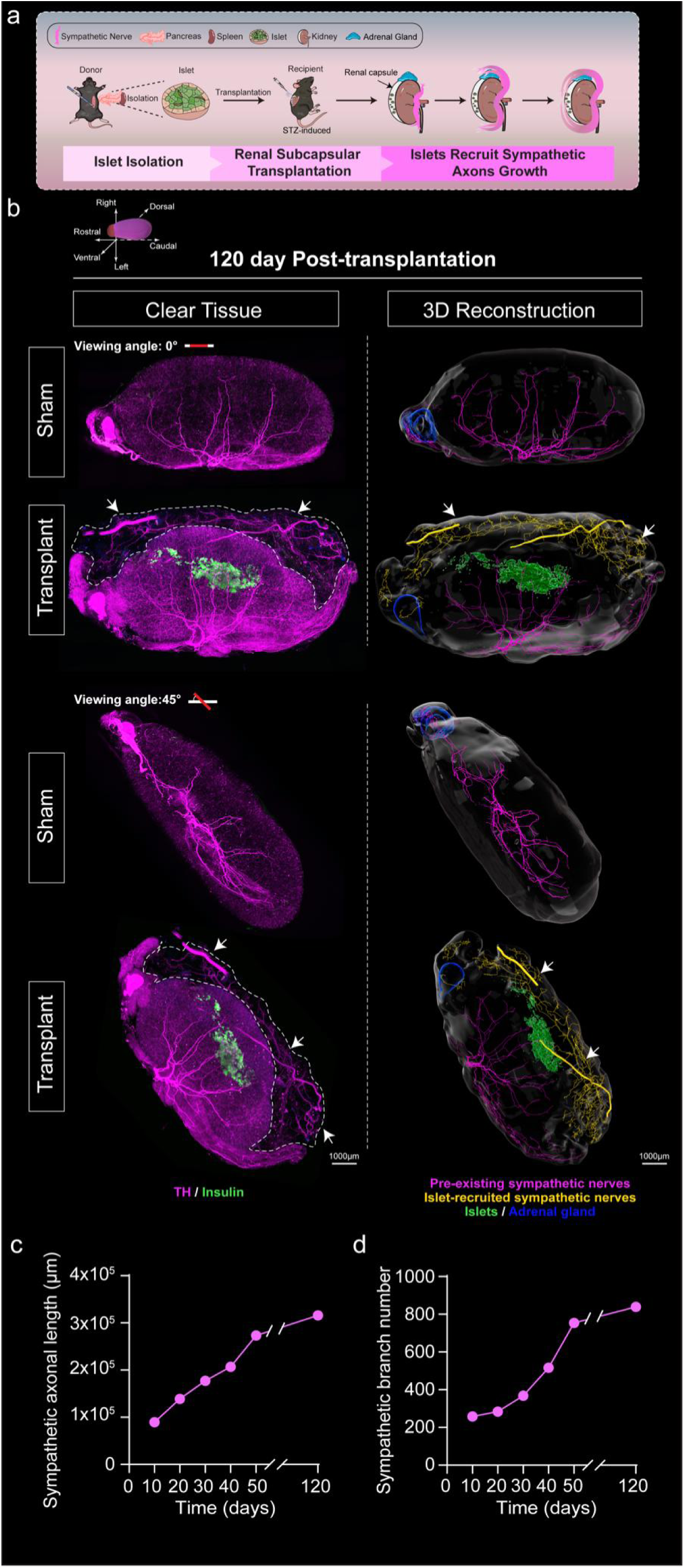
Pancreatic islets actively recruit sympathetic nerves independent of the pancreatic environment. **a,** Schematic illustration of the experimental strategy for renal subcapsular transplantation of approximately 500 isolated pancreatic islets into STZ-induced diabetic mice, to evaluate sympathetic nerve recruitment by islets transplanted outside their native pancreatic environment. **b,** Representative whole-kidney clearing images (left) showing sympathetic innervation (TH, magenta), transplanted islets (insulin, green) at day 120 post-transplantation. Corresponding 3D reconstructions (right) illustrate pre-existing sympathetic nerves within the kidney (magenta), islet-recruited sympathetic nerves (yellow), transplanted islets (green), and the adrenal gland (blue), viewed horizontally (0°) and from a tilted angle (45°). White arrowheads indicate sympathetic fibers clearly directed toward the transplanted islets. Sham-operated mice served as controls and exhibited no significant sympathetic nerve recruitment. Scale bars: 1000 µm. See Supplementary Video 4 for dynamic visualization. **c, d,** Quantitative analyses of sympathetic nerve growth recruited by transplanted islets, including total nerve length (c) and branch number (d), assessed at multiple time points (days 10, 20, 30, 40, 50, and 120) post-transplantation. Corresponding whole-tissue clearing and 3D reconstructions for each time point are presented in Supplementary Fig.3 and Supplementary Video 4-9.

**Figure 3.**
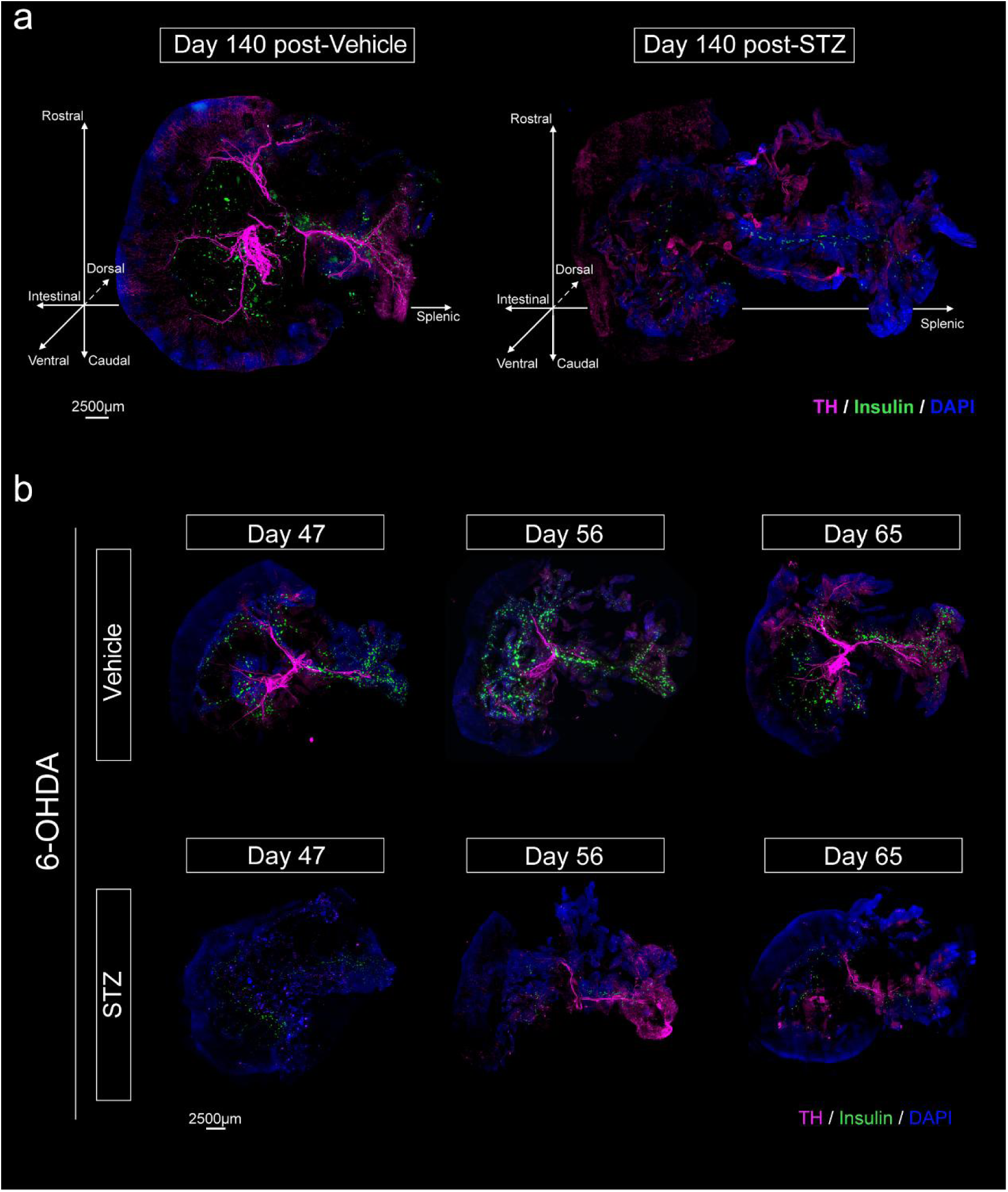
Islets maintain pancreatic sympathetic innervation and facilitate regeneration following denervation. **a,** Representative whole-pancreas 3D imaging showing sympathetic nerves (TH, magenta), pancreatic islets (insulin, green), and nuclei (DAPI, blue) in control mice (left) and mice subjected to chronic islet injury induced by STZ for 140 days (right). Scale bars: 2500 µm. See Supplementary Videos 10 and 11. **b,** Time-course imaging of sympathetic nerve regeneration following selective sympathetic nerve ablation using intrapancreatic 6-hydroxydopamine (6-OHDA) injection. Upper: significant sympathetic (TH, magenta) regeneration and restoration of structured associations with islets (insulin, green) at days 47, 56, and 65 post-denervation. Lower: impaired sympathetic regeneration at corresponding time points in mice subjected to severe islet injury induced by STZ administration one day post-denervation. Scale bars: 2500 µm. See Supplementary Videos 15-20.

We performed whole-kidney 3D imaging at day 120 post-transplantation, which clearly demonstrated sympathetic innervation of transplanted islets (islets, green; sympathetic nerves, magenta) from both horizontal (0°) and tilted (45°) perspectives (Fig. 2b; Supplementary Video 4). At day 120, sympathetic fibers were notably extended toward the transplanted islets (Fig. 2b, left; white arrowheads). To capture the dynamics of sympathetic nerve recruitment, we conducted tissue clearing imaging at intermediate time points (days 10, 20, 30, 40, and 50; Extended Data Fig.3a left; Supplementary Videos 5-9). Directional nerve growth toward transplanted islets was evident as early as day 10 and progressively strengthened. Quantitative analysis (Extended Data Fig.3c, d) revealed a significant increase in sympathetic nerve growth and branching at day 50 compared to day 10.

We performed detailed 3D reconstruction to clearly distinguish newly extended, islet-recruited sympathetic nerves (yellow, indicated by white arrowheads) from pre-existing renal sympathetic nerves (magenta), along with transplanted islets (green) and the adrenal gland (blue) (Fig. 2b, right; Extended Data Fig.3a, right; Supplementary Videos 4-9). Quantitative analysis revealed a progressive increase in sympathetic nerve length and branching toward transplanted islets, reaching significantly elevated levels by 120 days post-transplantation (Fig.2c, d).

### 3. Islets maintain sympathetic innervation and promote regeneration after denervation

Based on our finding that transplanted islets actively recruit sympathetic nerves in an ectopic environment, we next explored whether islets similarly regulate sympathetic innervation within their native pancreatic environment.

Initially, during experiments involving chronic islet injury induced by STZ, we unexpectedly observed pronounced disruption of pancreatic sympathetic nerves approximately 140 days post-treatment (Fig. 3a, right; Supplementary Video 10). In contrast, control mice receiving solvent injections maintained sympathetic nerve architecture (Fig. 3a, left; Supplementary Video 11). This finding revealed a previously unrecognized role of islets in maintaining pancreatic sympathetic nerve integrity.

Subsequently, to determine whether islets exhibit similar nerve-attracting properties within the native pancreas, we selectively ablated sympathetic nerves via intrapancreatic injection of 6-hydroxydopamine (6-OHDA). Successful denervation was confirmed at early time points (Extended Data Fig.4; Supplementary Videos 12-14). As anticipated, time-course analyses (days 47, 56, and 65 post-denervation) revealed progressive regeneration of sympathetic nerves and re-established organized interactions with islets (Fig. 3b, upper; Supplementary Videos 15-17). By contrast, severe islet injury induced by STZ administration one day post-denervation markedly impaired nerve regeneration, resulting in sparse, disorganized sympathetic fibers (Fig. 3b, lower; Supplementary Videos 18-20). Control mice injected with solvents maintained sympathetic innervation, ruling out nonspecific effects of vehicle injections. (Extended Data Fig.5; Supplementary Videos 21-23).

These results collectively revealed an essential and previously unrecognized role of pancreatic islets in maintaining sympathetic nerve integrity under physiological conditions and actively promoting nerve regeneration following injury, likely through the secretion of specific molecular factors.

### 4. Sympathetic denervation induces islet-specific upregulation of Reg-family proteins, especially Reg2 and Reg3β

To investigate the molecular mechanisms underlying islet-induced sympathetic nerve regeneration, we utilized the in situ sympathetic nerve ablation model induced by 6-OHDA described above. On day 7 post-denervation, pancreatic islets were directly isolated from their native pancreatic environment by laser-capture microdissection (LCM) (Fig. 4a). This approach preserved the native state of the islets and avoided potential alterations in their secretory profiles during isolation. Subsequent mRNA sequencing analysis revealed significant changes in gene expression, particularly a marked upregulation of regenerating gene family (Reg-family) members, notably *Reg2* and *Reg3b* (Fig. 4b).

**Figure 4.**
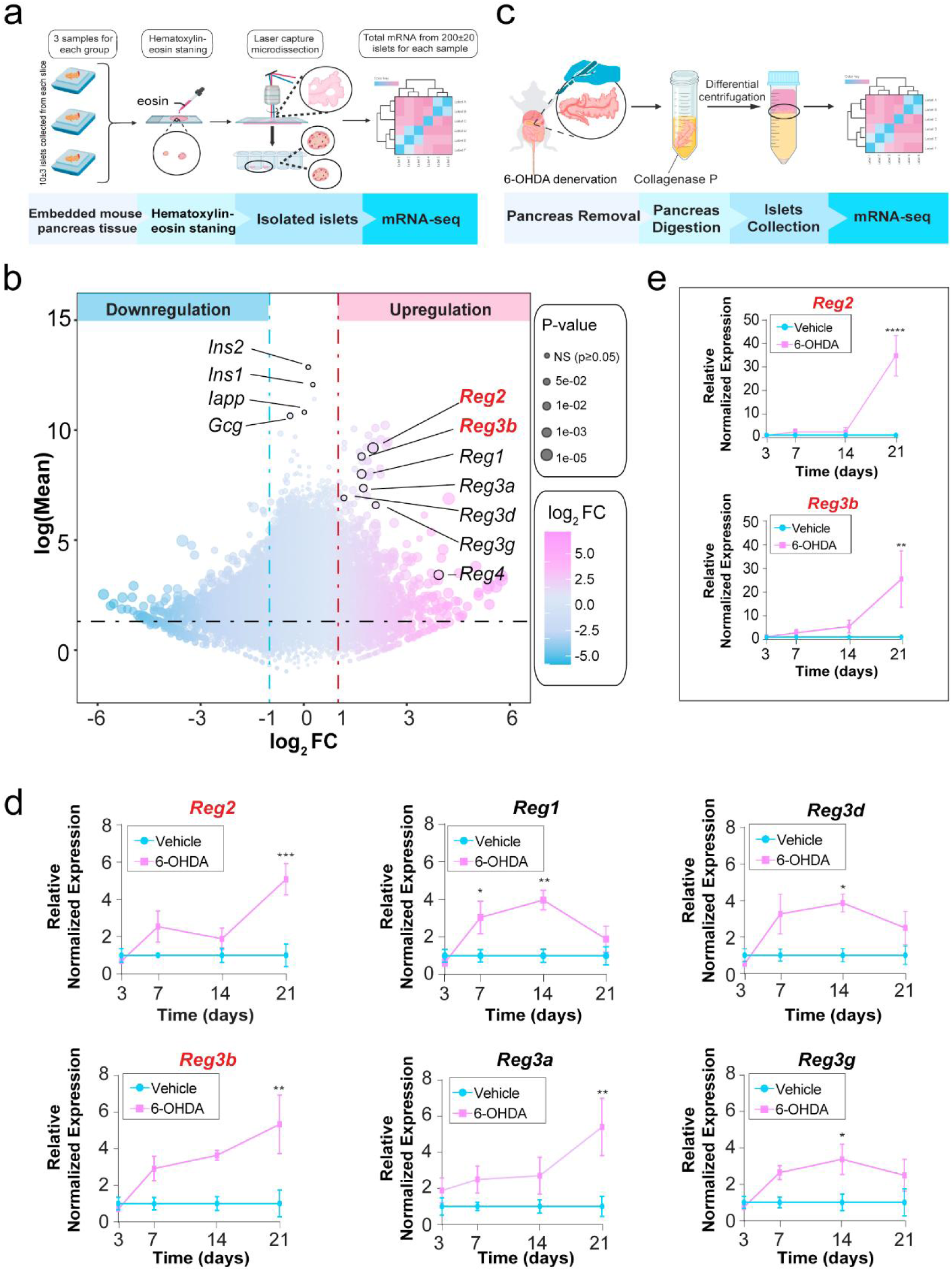
Reg-family proteins, particularly Reg2 and Reg3β, are prominently upregulated in pancreatic islets following sympathetic denervation. **a,** Schematic illustration of laser-capture microdissection (LCM) procedure for precise isolation of pancreatic islets from pancreas tissue on day 7 post-sympathetic denervation for mRNA sequencing. **b,** mRNA sequencing analysis of LCM-isolated islets revealed different expression genes (DEGs) post-denervation (n = 3 per group). **c,** Experimental workflow illustrating conventional isolation of pancreatic islets at multiple time points (days 3, 7, 14, and 21) post-sympathetic denervation for RNA sequencing analysis. **d,** mRNA sequencing-based temporal expression profiles of Reg-family genes in conventionally isolated islets post-denervation (n = 3 per group per time point). **e,** Real-time quantitative PCR (RT-qPCR) validation of *Reg2 and Reg3b* expression at indicated time points using samples from (d) (n = 3 per group per time point). Data presented as mean ± SEM, analyzed by two-tailed Student’s t-test. \**P* < 0.05, \*\**P* < 0.01, \*\*\**P* < 0.001.

To further clarify the dynamic expression patterns of Reg-family genes following sympathetic nerve injury, and to obtain sufficient islet samples for analysis at multiple time points, we subsequently utilized conventional islet isolation methods to collect islets at days 3, 7, 14, and 21 post-denervation for additional mRNA sequencing analysis (Fig. 4c). Consistent with the LCM-based findings, most Reg-family members, particularly *Reg2 and Reg3b*, exhibited significantly elevated expression levels from day 7 onward (Fig. 4d). These findings were independently validated by real-time quantitative PCR (RT-qPCR) analysis, confirming significant upregulation of *Reg2 and Reg3b* (Fig. 4e). Given that Reg-family proteins are well-established secretory factors implicated in tissue repair and regeneration^32–34^, our results strongly suggest that islet-derived Reg-family proteins, particularly Reg2 and Reg3β, serve as key molecular cues guiding sympathetic nerve regeneration toward pancreatic islets.

### 5. Exogenous Reg2 and Reg3β accelerate sympathetic nerve growth and enhance islet graft function

To functionally validate the importance of Reg2 and Reg3β and explore potential translational implications, we initially transplanted pancreatic islets encapsulated in a hydrogel matrix containing exogenous recombinant Reg2 (rReg2) and Reg3β (rReg3β) beneath the renal capsule of recipient mice. At day 50 post-transplantation, whole-kidney clearing and 3D imaging revealed that local administration of rReg2 and rReg3β significantly enhanced sympathetic nerve regeneration, as evidenced by increased nerve length, branching, and directional growth toward transplanted islets compared with controls (Fig. 5a, b; Supplementary Video 24).

**Figure 5.**
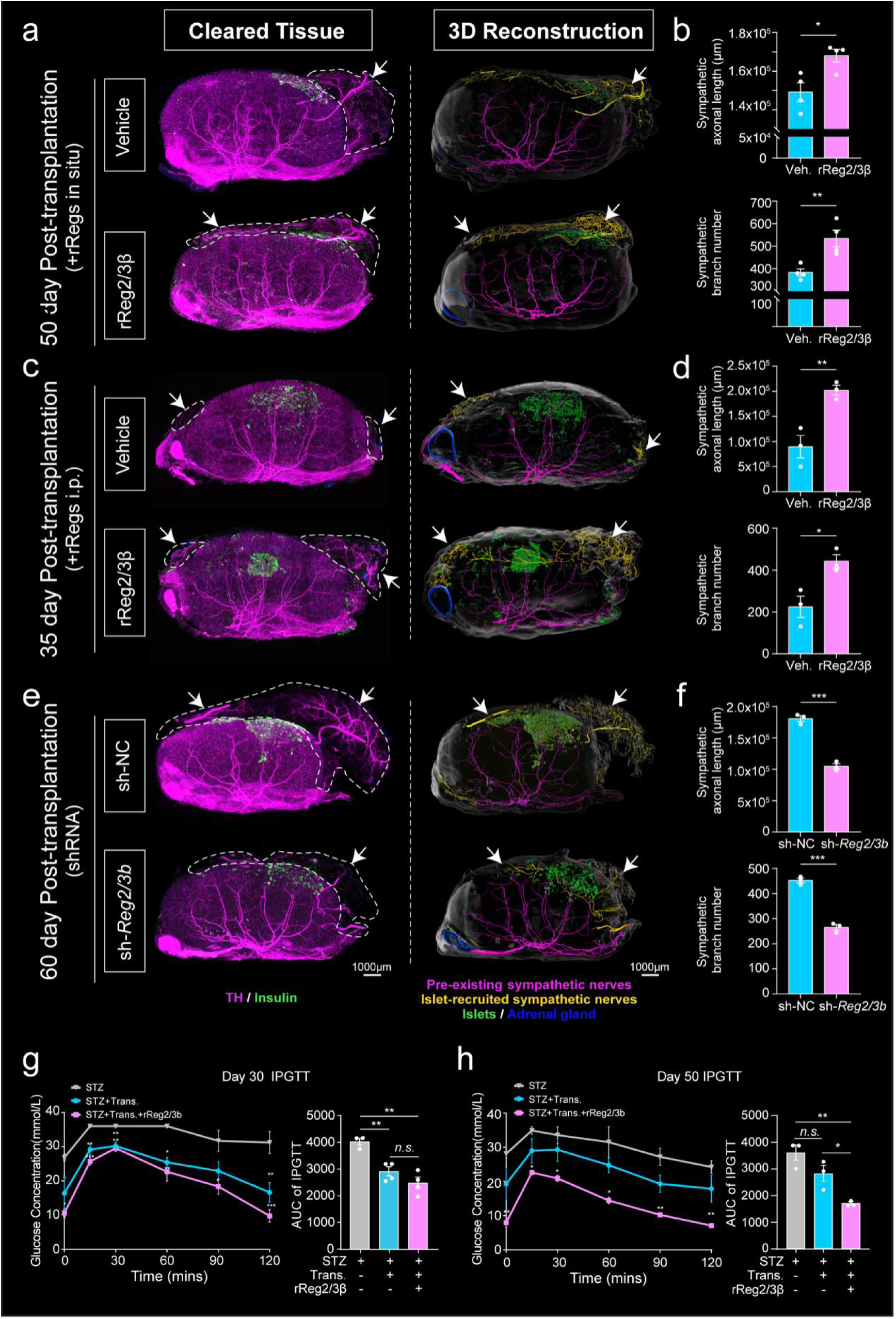
Exogenous rReg2 and rReg3β enhance sympathetic nerve regeneration and improve functional outcomes in transplanted islets. **a,** Representative whole-kidney clearing images (left) showing sympathetic nerves (TH, magenta), transplanted islets (insulin, green) at day 50 post-transplantation. Corresponding 3D reconstructions (right) illustrate pre-existing renal sympathetic nerves (magenta), islet-recruited sympathetic nerves (yellow), transplanted islets (green), and the adrenal gland (blue). White arrowheads indicate sympathetic nerves extending toward islets. Recombinant regenerating protein 2 (rReg2) and recombinant regenerating protein 3 beta (rReg3β) proteins were incorporated into hydrogel during gelation, thereby achieving localized drug delivery to islet grafts. Scale bar, 1000 μm. See Supplementary Video 24. **b,** Quantification of sympathetic nerve fibers extending toward transplanted islets following localized hydrogel delivery of rReg2 and rReg3β proteins, as illustrated in (a) (n = 4 per group). **c,** Representative whole-kidney clearing images (left) at day 35 post-transplantation showing sympathetic nerves (TH, magenta), transplanted islets (insulin, green). Corresponding 3D reconstructions (right) illustrate pre-existing renal sympathetic nerves (magenta), islet-recruited sympathetic nerves (yellow), transplanted islets (green), and the adrenal gland (blue). White arrowheads indicate sympathetic nerves extending toward islets. rReg2 and rReg3β proteins were systemically administered via intraperitoneal (i.p.) injections on days 1, 3, and 5 post-transplantation. Scale bar, 1000 μm. See Supplementary Video 25. **d,** Quantification of sympathetic nerve fibers extending toward transplanted islets following i.p. injection of rReg2 and rReg3β proteins, as illustrated in (c) (n = 3 per group). **e,** Representative whole-kidney clearing images (left) at day 60 post-transplantation showing sympathetic nerves (TH, magenta) and transplanted islets (insulin, green). Corresponding 3D reconstructions (right) illustrate pre-existing renal sympathetic nerves (magenta), islet-recruited sympathetic nerves (yellow), transplanted islets (green), and adrenal gland (blue), following lentiviral-mediated knockdown of *Reg2* and *Reg3b*. White arrowheads indicate sympathetic nerves extending toward islets. Knockdown efficiency was verified by RT-qPCR (Supplementary Fig.5d); shRNA sequences provided in Supplementary Table 1. Scale bar, 1000 μm. See Supplementary Video 26. **f,** Quantification of sympathetic nerve fibers extending toward transplanted islets following lentiviral-mediated knockdown of *Reg2* and *Reg3b*, as illustrated in (e) (n = 3 per group). **g, h,** IPGTT at days 30 (g) and 50 (h) post-transplantation. Notably, the enhancement in islet graft function was more pronounced at day 50 than at day 30 (n = 3-4 per group per time point). Corresponding whole-kidney clearing images are shown in Supplementary Video 27. Data are shown as mean ± SEM, analyzed by two-tailed Student’s t-test. \**P* < 0.05, \*\**P* < 0.01, \*\*\**P* < 0.001.

Given the potential limitations of local protein delivery in clinical contexts, we next assessed the efficacy of a more practical route — intraperitoneal administration of rReg2 and rReg3β — in promoting sympathetic regeneration. Recombinant proteins were administered intraperitoneally (i.p.) on days 1, 3, and 5 post-transplantation. Tissues analyzed 35 days post-transplantation showed that systemic delivery also robustly promoted sympathetic regeneration, with significantly increased nerve length and branching toward transplanted islets (Fig. 5c, d; Supplementary Video 25).

To further confirm that endogenous Reg2 and Reg3β specifically mediate sympathetic nerve regeneration toward islets, we used lentiviral-mediated RNA interference to knock down *Reg2* and *Reg3b* expression in isolated islets prior to transplantation (lentiviral sequences provided in Supplementary Table 1), and knockdown efficiency was confirmed by RT-qPCR analysis (Extended Data Fig.6d). At day 60 post-transplantation, sympathetic regeneration toward islets with *Reg2* and *Reg3b* knockdown was significantly delayed, characterized by reduced nerve length and branching compared to controls (Fig. 5e, f; Supplementary Video 26).

Representative whole-kidney clearing images and corresponding 3D reconstructions (horizontal 0° view, Fig. 5a, c, e; titled 45° view, Extended Data Fig.6a-c) comprehensively illustrate the spatial interactions between sympathetic nerves and transplanted islets under these experimental conditions. White arrowheads indicate sympathetic fibers clearly directed toward the transplanted islets.

To evaluate whether Reg2 and Reg3β-mediated sympathetic nerve regeneration functionally enhances transplanted islets, we administered combined local and systemic rReg2 and rReg3β treatment during islet transplantation. Subsequently, intraperitoneal glucose tolerance tests (IPGTT) were performed at days 30 and 50 post-transplantation. Mice receiving rReg2 and rReg3β-treated islets exhibited significantly improved glucose tolerance compared to controls receiving islets alone. Notably, this functional enhancement was more pronounced at day 50 than at day 30 (Fig. 5g, h). These results suggest that Reg2 and Reg3β not only accelerate but also support prolonged functional performance of transplanted islets, highlighting the therapeutic promise of Reg proteins for enhancing clinical outcomes in islet transplantation and managing diabetes more effectively (Supplementary Video 27).

## Discussion

Pancreatic islets, essential regulators of glucose homeostasis, are dispersed micro-organs within the pancreas^35,36^. This unique distribution enables islets to precisely monitor and rapidly respond to glucose fluctuations, yet raises a fundamental physiological question: how do dispersed islets achieve synchronized neural integration during acute metabolic stress, such as severe hypoglycemia or fight-or-flight responses^37–39^? Here, we reveal for the first time a precise, “wiring-light bulb” anatomical arrangement, where islets (“light bulbs”) align closely along fine sympathetic nerve branches (“wiring”). This specialized organization provides a structural foundation for rapid and coordinated neural integration, analogous to multiple bulbs simultaneously illuminating once electrical current passes through the wiring, highlighting how “**structure dictates function”** in neural signaling.

Using renal subcapsular transplantation, we confirmed that pancreatic islets actively shape their specialized sympathetic innervation independently of the pancreatic environment, and similarly promote sympathetic nerve regeneration following injury within the native pancreas. Notably, chronic islet injury resulted in progressive disruption of pancreatic sympathetic networks, further uncovering a previously unrecognized role of islets in actively maintaining sympathetic neural integrity. These findings collectively extend the classical view of endocrine organs beyond passive neural targets^40,41^, revealing instead their active participation in both establishing and preserving neural architecture. This introduces a novel biological paradigm for organ-nerve interactions: **function shapes and sustains structure**, a principle potentially applicable to other endocrine organs.

Here, we identified Reg-family proteins, particularly Reg2 and Reg3β, as critical mediators enhancing both islet graft function and sympathetic nerve regeneration, with therapeutic efficacy notably increasing over time. Traditionally associated with inflammation^42,43^, tissue repair^44^, and regeneration^34,45^, Reg proteins had not previously been studied in a neuro-endocrine context. Our findings substantially expand the functional repertoire of these proteins, uncovering their novel capacity to directly facilitate sympathetic nerve growth and promote the functional maturation of transplanted islets. This dual role positions Reg2 and Reg3β as promising therapeutic candidates not only for improving clinical outcomes in islet transplantation but also for preserving sympathetic nerve integrity compromised during diabetes progression.

Further exploration of their translational potential could provide innovative strategies for managing metabolic and neural complications associated with diabetes.

Despite these advances, key questions remain unanswered. First, it remains unclear whether the specialized anatomical arrangement between sympathetic nerves and pancreatic islets observed in mice is conserved in humans. Second, although we identified Reg2 and Reg3β as critical mediators of sympathetic nerve regeneration, their specific cellular sources, molecular mechanisms, and signaling pathways within islets require further elucidation. Future studies using human islets or organoids combined with single-cell RNA sequencing and spatial transcriptomics could address these gaps. Finally, optimal delivery methods, precise dosage, and combinations with other neurotrophic factors must be defined to facilitate clinical translation.

In summary, our findings redefine the fundamental relationship between endocrine organs and neural networks, uncovering a previously unrecognized biological principle whereby pancreatic islets actively construct and sustain their specialized sympathetic architecture. This reveals an innovative paradigm in neuro-endocrine biology: endocrine organs are not passively regulated by neural inputs but instead actively shape and maintain neural circuits to optimize physiological outcomes. By identifying Reg2 and Reg3β as critical molecular orchestrators, we open avenues for novel therapeutic strategies not only for diabetes but potentially for broader metabolic and neurological disorders where neural integrity is compromised. Future research should explore whether similar organ-driven neural remodeling occurs across other endocrine systems, providing generalizable insights into organ-nerve interactions and expanding therapeutic possibilities for metabolic and neurological disorders.

## Methods

### Ethical approval

All animal experiments conducted in this study were approved by the Institutional Animal Ethics Committee of the Shanghai Cancer Institute (A2024170-004), Ren Ji Hospital, School of Medicine, Shanghai Jiao Tong University. Experiments were performed strictly in accordance with institutional guidelines and ethical standards.

### Animals

Male C57BL/6J mice (8-10 weeks old) were obtained from Vitalriver and maintained under specific pathogen-free conditions (temperature: 22 ± 2 °C; humidity: 55% ± 5%; 12-h light/dark cycle) at Shanghai Jiao Tong University, with ad libitum access to standard rodent chow and water. Body weight, food intake, and water consumption were recorded regularly. All animal procedures were approved by the Institutional Animal Care and Use Committee (IACUC) of Shanghai Jiao Tong University and followed the NIH Guide for the Care and Use of Laboratory Animals.

### Whole-organ clearing and labeling

#### Tissue harvesting

Mice were deeply anesthetized via intraperitoneal (i.p.) injection of sodium pentobarbital (50 mg/kg, RWD, r510-22) and perfused transcardially with phosphate-buffered saline (PBS, Servicebio, G4202) followed by 4% paraformaldehyde (PFA, Servicebio, G1101). The pancreas was carefully dissected, rinsed in PBS, and placed on a moist filter paper. To preserve the organ’s spatial architecture, the pancreas was gently spread out and pinned at the edges using fine needles (Extended Data Fig.1a).

#### 3D immunolabeling

The sample was treated with methanol solutions for 3 days at room temperature and bleached with a 5% H₂ O₂ solution overnight at 4°C. Then, the sample was treated with 0.2% Triton X-100 solution for 3 days at room temperature with shaking. Subsequently, the sample was immersed in a 10% donkey serum blocking solution for 3 days at room temperature, followed by incubation in the primary antibody solution (TH, 1:500, rabbit, ab112, Abcam; Insulin, 1:500, rat, MAB1417, R&D Systems) for 6 days at room temperature, and then in the secondary antibody solution (donkey anti-rabbit 561 nm, 1:500, A32794, Invitrogen; donkey anti-rat 488 nm, 1:500, A21208, Invitrogen; DAPI, 1:500, D1306, Invitrogen) for 6 days at room temperature. After immersion in the antibody solutions, the sample was washed with 0.2% Tween-20 solution for 2 days at room temperature^46^. Detailed antibody information is summarized in Table 1.

### Clearing method

The FDISCO (fluorescence imaging of solvent-cleared organs with superior fluorescence-preserving capability) tissue clearing reagent kit (JA11012, Jarvisbio) was used in this study^47^. The clearing procedure was performed strictly according to the manufacturer’s instructions. Briefly, labeled mouse pancreas samples were sequentially incubated in reagents A, B, C, and D for 6 hours each, followed by reagent E for 12 hours, and finally incubated in reagent F until complete transparency was achieved. All incubations were conducted at 6-8 °C with gentle shaking.

### 3D fluorescence imaging and digital reconstruction of cleared tissues

#### Light-sheet microscope

The cleared samples were imaged using a light sheet microscope (LiToneXL, Light Innovation Technology) equipped with a 4 × objective lens (NA = 0.28, working distance = 20 mm). Thin light sheets were illuminated from the four sides of samples, and a merged image was saved. To acquire images, the cleared samples were manually attached to the sample holder adapter. Subsequently, the samples were immersed in imaging reagent within a 3D printing sample chamber and excited with light sheets of different wavelengths.

### Confocal microscope

The cleared tissue samples were mounted on two cover glasses and imaged using an inverted confocal fluorescence microscope (LSM880, Zeiss, Germany) equipped with the Plan Apochromat 10×/0.45 objective (dry lens; working distance = 2.0 mm), Plan Apochromat 20 × /0.8 objective (dry lens; working distance = 0.55 mm), and Plan Apochromat 40×/0.95 objective (water immersion lens; working distance = 0.20 mm). The collected images were subsequently processed with Zen software (Zeiss, Germany) and reconstructed into 3D images using Imaris software (Version 7.2.3, Bitplane, Switzerland)^48^.

### Imaging data processing

The resulting TIFF raw data images were stitched and converted using LitScan software and subsequently analyzed with Imaris software^48^.

### Digital reconstruction and quantitative analysis of cleared nerve fibers

The Filament module of Imaris software was used for 3D tracing of nerve fibers in cleared tissues. Cone mode and AutoDepth mode were selected to accurately locate and delineate the nerve fibers, and the segment diameters were adjusted according to the actual diameters of nerve trunks and branches. Finally, filament length and branching points were quantified using the statistics panel of the Filament module.

### Islet isolation and renal subcapsular transplantation

Pancreatic islets were isolated from donor mice using collagenase P (Roche, CollP-RO) digestion and density gradient centrifugation as previously described^49^. Approximately 500 islets were evenly divided into two parts and transplanted under the bilateral renal capsule of STZ-induced diabetic recipient mice. For transplantation, mice were anesthetized, and a small incision was made to expose the kidney. Islets were transferred using a Hamilton syringe and gently injected under the renal capsule. The incision was closed with surgical sutures, and mice were allowed to recover with appropriate postoperative care.

### Establishment of diabetic mice models

Two distinct protocols for streptozotocin (STZ)-induced diabetes were employed, depending on the experimental purpose. Specifically, a multiple low-dose STZ protocol was used to establish stable diabetes for subsequent islet transplantation experiments. A single high-dose STZ injection was administered one day after sympathetic nerve ablation using 6-hydroxydopamine (6-OHDA) to rapidly induce severe β-cell injury, thus preventing potential confounding effects of residual functional islets on sympathetic nerve regeneration.

### Multiple low-dose STZ induction

Diabetes was induced by daily i.p. injections of STZ (50 mg/kg) dissolved in 0.1 M citrate buffer (pH 4.2-4.5), administered over five consecutive days^50^.

### Single high-dose STZ induction following 6-OHDA treatment

Diabetes was induced by a single high-dose i.p. injection of STZ (150 mg/kg), also dissolved in 0.1 M citrate buffer (pH 4.2-4.5)^51^. The injection was given following an 8-hour fasting period, specifically one day after pancreatic sympathetic nerve ablation using 6-OHDA.

Blood glucose levels were monitored every other day using a glucometer (Accu-Chek, Roche). Mice were defined as diabetic when blood glucose exceeded 16.7 mmol/L in three consecutive measurements.

### Blood glucose measurement and IPGTT

Random blood glucose levels were measured from tail vein blood using a glucometer. IPGTT were performed after an 8-hour fasting. Mice received an i.p. injection of glucose (2 g/kg), and blood glucose levels were measured at 0, 15, 30, 60, and 120 min post-injection^52^.

### Sympathetic nerve ablation

10-week-old mice were randomly assigned into the vehicle group and the 6-OHDA group. After anesthetizing the mice with isoflurane, the hair in the abdomen was removed and the area was disinfected with povidone-iodine. Then, a small incision was made along the midline of the abdomen skin to expose the pancreas. The pancreas was pulled out gently and fully exposed with sterile tweezers. 6-OHDA (HY-B1081, MCE) was administered by intrapancreatic injection (1 mg in 15 µL saline containing 0.1% L-ascorbic acid [HY-B0166, MCE], 100 mg/kg). To pharmacologically ablate sympathetic nerve in the whole pancreas, this 15-μL dose of 6-OHDA was evenly administered from the pancreas head to the tail. Control mice received a 15-μL dose of 0.1% L-ascorbic acid injections.

### Regenerating protein family (Reg-family) protein administration

Recipient mice were randomly divided into four experimental groups:

#### Local administration

Recombinant Reg2 (rReg2) and recombinant Reg3β (rReg3β) proteins (500 nM final concentration) were incorporated into a biocompatible hydrogel matrix (Corning, Cat#354236) at a final concentration of 2 mg/mL. Isolated islets were gently mixed with the Reg-loaded hydrogel. Controls received islets embedded in hydrogel without Reg proteins.

### Systemic administration

Mice received purified islets embedded in a hydrogel matrix (2 mg/mL, Corning, Cat#354236). For the systemic administration group, mice additionally received i.p. injections of rReg2 and rReg3β protein mixture (150 mg/kg, dissolved in sterile ddH_2_O at 100 mg/mL) on days 1, 3, and 5 post-transplantation^53,54^, starting on the transplantation day. Controls received equal volumes of sterile ddH_2_O on the same schedule.

All animal procedures were performed under general anesthesia, following institutional animal care and use guidelines.

### Lentiviral-mediated knockdown of *Reg2* and *Reg3b* expression in pancreatic islets

#### shRNA design and vector construction

Three independent shRNA sequences targeting mouse *Reg2* (NCBI Gene ID: NM_009043.2) and *Reg3b* (NCBI Gene ID: NM_011036.1), along with a scrambled control shRNA, were designed and synthesized **(Supplementary Table 2)**. Each sequence was cloned into the lentiviral vector pSLenti-U6-shRNA-CMV-EGFP-F2A-Puro-WPRE.

### Validation of shRNA knockdown efficiency in isolated islets

Freshly isolated mouse pancreatic islets (approximately 100 islet equivalents, IEQ) were pre-cultured overnight in RPMI-1640 medium supplemented with 10% fetal bovine serum (FBS). Equal volumes of lentiviral preparations containing three distinct shRNAs targeting either *Reg2* or *Reg3b* were pre-mixed at a 1:1:1 ratio, ensuring equal multiplicity of infection (MOI) distribution for each shRNA construct.

The pooled lentiviral mixtures were added to the pre-cultured islet at a total MOI of 10-20 in the presence of polybrene (10 μg/mL). After incubation at 37°C for 8 h, the virus-containing medium was replaced with fresh culture medium. Gene knockdown efficiency was confirmed by RT-qPCR analysis prior to islet transplantation (see Extended Data Fig.7).

### Lentiviral-mediated shRNA delivery during islet transplantation

To achieve efficient *Reg2* and *Reg3b* knockdown during islet transplantation, freshly isolated pancreatic islets were mixed with lentiviral preparations (containing the same shRNA mixture described above) and rapidly encapsulated in a hydrogel matrix. The islet-lentivirus-hydrogel mixture was then incubated for approximately 30 min at 37°C to allow complete gelation, and subsequently transplanted beneath the renal capsule of anesthetized recipient mice.

### Real-time quantitative PCR (RT-qPCR)

Total RNA was extracted from tissues or cells using TRIzol reagent (Invitrogen). cDNA was synthesized using the PrimeScript RT reagent Kit (Takara). RT-qPCR was performed with SYBR Green PCR Master Mix (Applied Biosystems) on a QuantStudio 6 Flex Real-Time PCR System (Applied Biosystems). Gene expression levels were normalized to β-actin and calculated using the 2^^-ΔΔCt^ method. Primer sequences are listed in Supplementary Table 3.

### Transcriptome sequencing of laser-captured pancreatic islets

#### Sample preparing

Formalin-fixed, paraffin-embedded (FFPE) pancreatic tissue was sectioned at 7 μm thickness. Slides were first deparaffinized in xylene for 5 min, rehydrated sequentially through graded ethanol solutions (100%, 95%, 70%, and 50%; 2 min each), and rinsed briefly in distilled water. Sections were then rapidly stained with hematoxylin and eosin (H&E) using an optimized protocol designed to preserve RNA integrity. Following staining, slides were dehydrated in ascending ethanol concentrations, cleared with xylene, and allowed to air-dry completely under RNase-free conditions before further processing.

### Laser-capture microdissection (LCM) and libraries preparation

LCM was performed using a PALM MicroBeam system (Zeiss MicroImaging, Munich, Germany). Islet regions were identified based on morphological criteria under microscopic observation. Approximately 300 islets per sample were precisely isolated via laser microdissection and collected into 200 μL adhesive-capped tubes (Carl Zeiss, cat.no. 415190-9191-000, Jena, Germany). Immediately following collection, 10 μL of LCM lysis mix buffer was added, and tubes were sealed tightly with Parafilm.

Transcriptome libraries were generated following the Smart-3SEQ protocol. Briefly, cell lysis and RNA fragmentation were performed at 60°C for 60 min, followed by the addition of 10 μL LCM TS-RT mix for reverse transcription. Amplification of the resulting cDNA and subsequent cleanup were performed according to the published protocol. The final library concentration was measured using a Qubit 3.0 fluorometer (Thermo Fisher Scientific, MA, USA)^55,56^.

### Next-generation sequencing

The cDNA libraries were profiled for size distribution on an Agilent 2100 with High Sensitivity D1000 reagent kit (Agilent, Santa Clara, USA) and then stored at −20°C until sequencing. The libraries were sequenced using the Illumina high-throughput sequencing platform NovaSeq 6000 system (Illumina, San Diego, USA) with 2×150 bp pair-end mode.

### Data analysis

The FASTQ data were analyzed according to the Smart-3SEQ protocol. Briefly, base calls from the NovaSeq were demultiplexed and converted to FASTQ format and adapter trimming were performed. The first 5nt of each read were removed from its sequence and appended to the read name as its UMI; the following 3nt were assumed to derive from the G-overhand of the template-switch oligonucleotide and were discarded. Reads were aligned by STAR to the mouse reference genome mm10. PCR duplicate reads that start at the same genome position with the same UMI sequence were detected and discarded. Then the transcript abundance was quantified. Differential gene expression was analyzed with DESeq2. The regularized log transformation was used to normalize read counts. PCA was performed on the normalized data with the prcomp function.

### Bulk RNA-seq of isolated pancreatic islets

Pancreatic islets were isolated from 0.1% L-ascorbic acid or 6-OHDA treated mice using collagenase P (Roche, CollP-RO) digestion and density gradient centrifugation as previously described. Total RNA was extracted from islet samples with the Universal RNA Extraction CZ Kit (RNC643, ONREW) according to the manufacturer’s instructions. RNA quantity was analysed using Qubit 4.0 (Invitrogen) and quality examined by electrophoresis on a denaturing agarose gel. RNA libraries were prepared using the VVAHTS® Universal V8 RNA-seq Library Prep Kit for Illumina (NR605-0, Vazyme), followed sequencing using the Illumina NovaSeq 6000 platform with the 150 paired-end sequencing strategy. Enrichment of mRNA, library construction, sequencing and data analysis were performed by Shanghai Xu Ran Biotechnology Co., Ltd. The raw data was handled by Skewer v0.2.2 (https://sourceforge.net/projects/skewer/files/?source=navbar) and data quality was checked by FastQC v0.11.2 (http://www.bioinformatics.babraham.ac.uk/projects/fastqc/). The read length was 2×150 bp. Clean reads were aligned to the mouse genome GRCm39 (mm10) from ensembl using STAR (https://github.com/alexdobin/STAR), with one mismatch allowed. StringTie (v1.3.1c) was used to generate gene expression data and differential gene expression was analysed by DESeq2 (v1.16.1) (https://bioconductor.org/packages/release/bioc/html/DESeq2.html). The thresholds for determining DEGs are *P* < 0.05 and absolute fold change ≥ 2. Then DEGs were chosen for function and signaling pathway enrichment analysis using TopGO (https://www.bioconductor.org/packages/release/bioc/html/topGO.html) and KEGG database (https://www.genome.jp/kegg/pathway.html). The significantly enriched pathways were determined when *P* < 0.05.

## Data availability

The raw sequencing data from the laser capture microdissected samples and the isolated islet samples have been deposited in the NCBI SRA under accession number PRJNA1241420 and PRJNA1243594 respectively. All other data in this article are available from the corresponding author on reasonable request. The datasets generated and analyzed during this study are available via Figshare (https://figshare.com/s/37e36235c0efd54a05a7).

## Acknowledgements

This study was supported by the National Natural Science Foundation of China (92168111, 82230087, 82350123, 82203228), the Shanghai Municipal Education Commission-Gaofeng Clinical Medicine Grant Support (20181708), Innovative research team of high-level local universities in Shanghai (SHSMU-ZDCX20210802), Shanghai Pilot Program for Basic Research - Shanghai Jiao Tong University (21TQ1400225), 111 project (NO. B21024), Shenyang Science and Technology Plan in 2022 (22-101-0-22). We thank Jarvis (Wuhan) Biological Pharmaceutical Co., Ltd. for technical support in tissue clearing and 3D imaging.

## Author contributions

Z.-G.Z. and D.-X.L. conceived the project and designed experiments. J.-M.L., J.-J.W., Y.-Z.Q., and D.-X.L. performed experiments and analyzed the data. D.-X.L. and J.-M.L. wrote the manuscript. J.-J.W. and H.-L. performed bioinformatics analyses. D.A., T.Y., Y.-X.H., W.-T.S., X.-Y.M., and M.M. assisted with experiments. L.Z., X.-M.Y., Y.Q., Y.-L.Z., and Y.-Q.Y. provided technical support. S.-H.J., J.L., L.-P.H., X.-L.Z., and Q.L. provided critical insights and revised the manuscript. All authors reviewed and approved the final manuscript.

