## supplementary materials for "A neuroendocrine principle: Pancreatic islets actively shape sympathetic innervation"

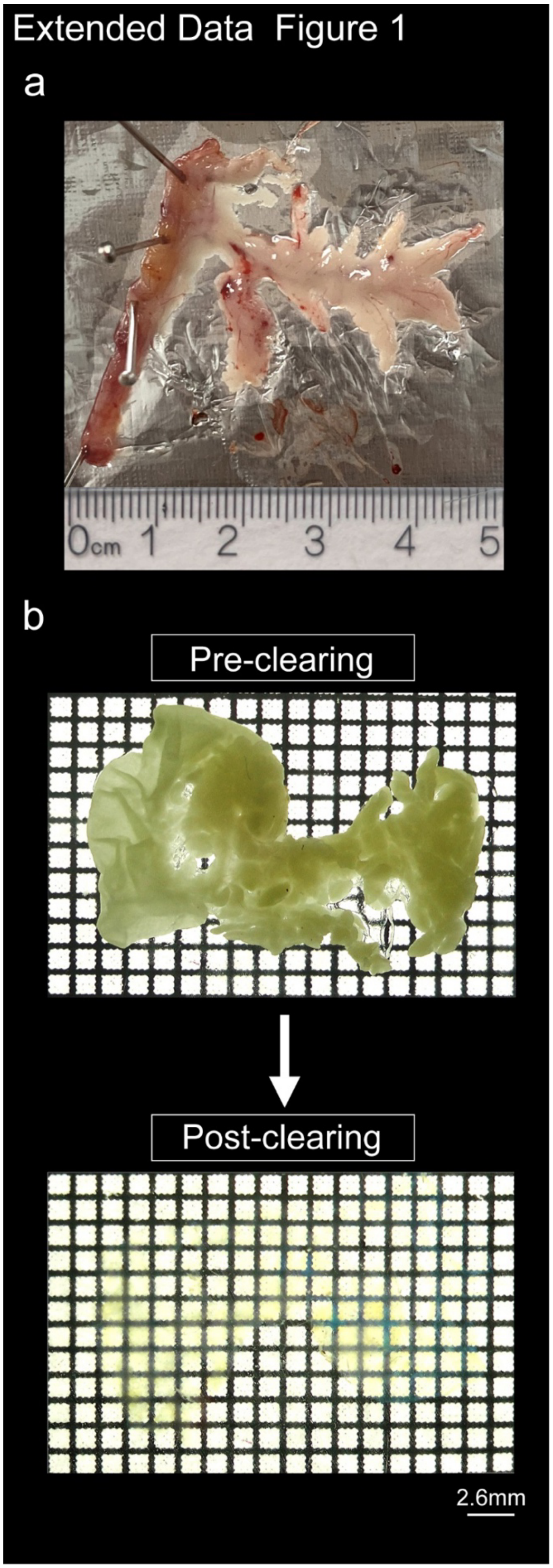
Extended Data Figure 1
a
0cm 1 2 3 4 5
b
Pre-clearing
Post-clearing
2.6mm

**Extended Data Figure 1. Methodology of pancreas unfolding and tissue clearing.**

**a,** Representative image demonstrating complete unfolding of the intact mouse pancreas, preserving its native spatial architecture.

**b,** Images of the unfolded mouse pancreas before and after whole-organ tissue clearing, illustrating increased tissue transparency.

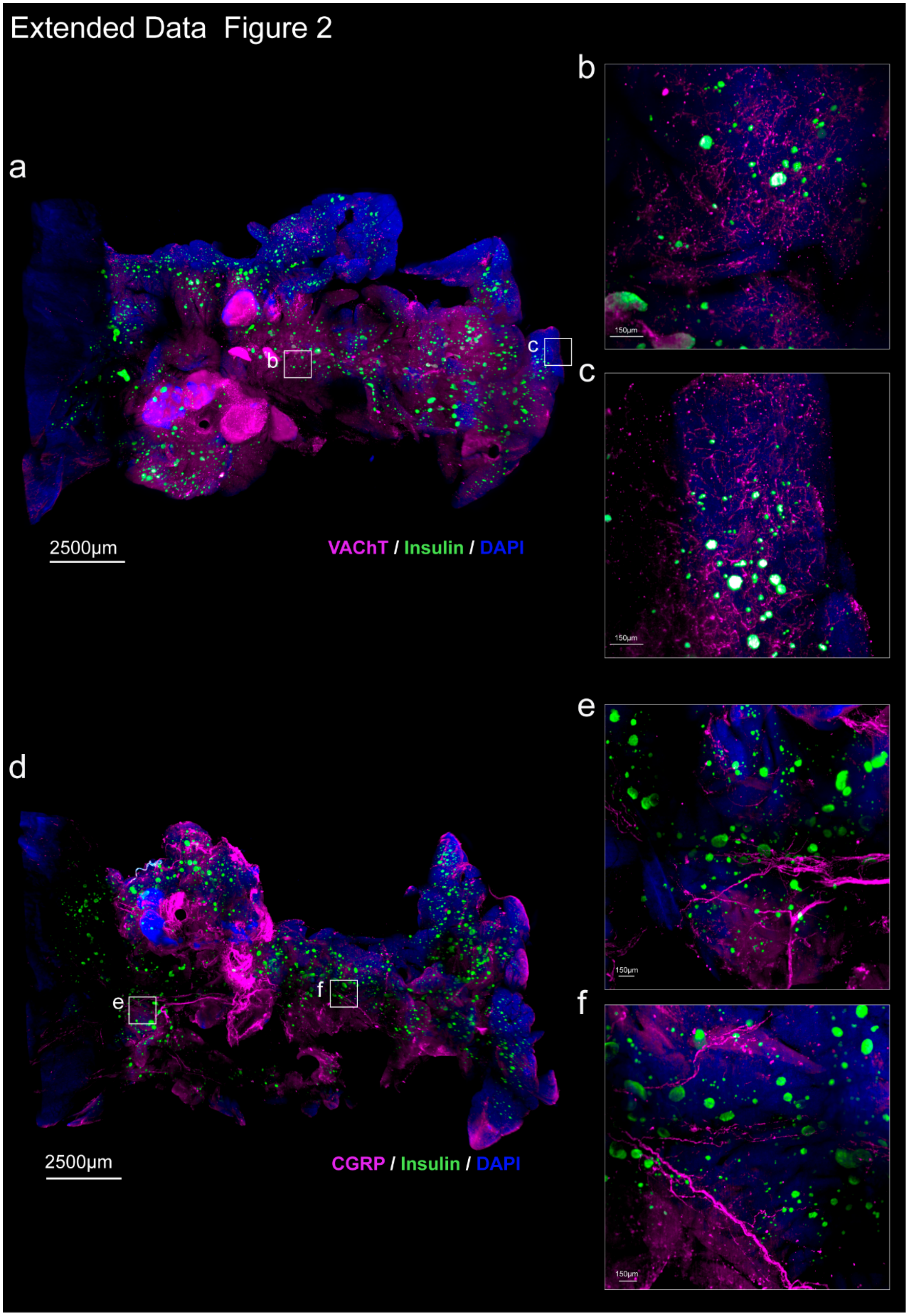
Extended Data Figure 2
a
b
c
2500µm
VAChT / Insulin / DAPI
150µm
d
e
f
2500µm
CGRP / Insulin / DAPI
150µm

**Extended Data Figure 2. Spatial relationship between pancreatic islets and parasympathetic or sensory nerves.**

**a,** Whole-pancreas 3D imaging of parasympathetic nerves (VAChT, magenta), islets (insulin, green) and nuclei (DAPI, blue). Scale bar: 2500 μm. See Supplementary Video 2.

**b, c,** Magnified views of selected regions from (a). Scale bars: 150 μm.

**d,** Whole-pancreas 3D imaging of sensory nerves labeled with CGRP (magenta). Scale bar: 2500 μm. See Supplementary Video 3.

**e, f,** Magnified views of selected regions from (d). Scale bars: 150 μm.

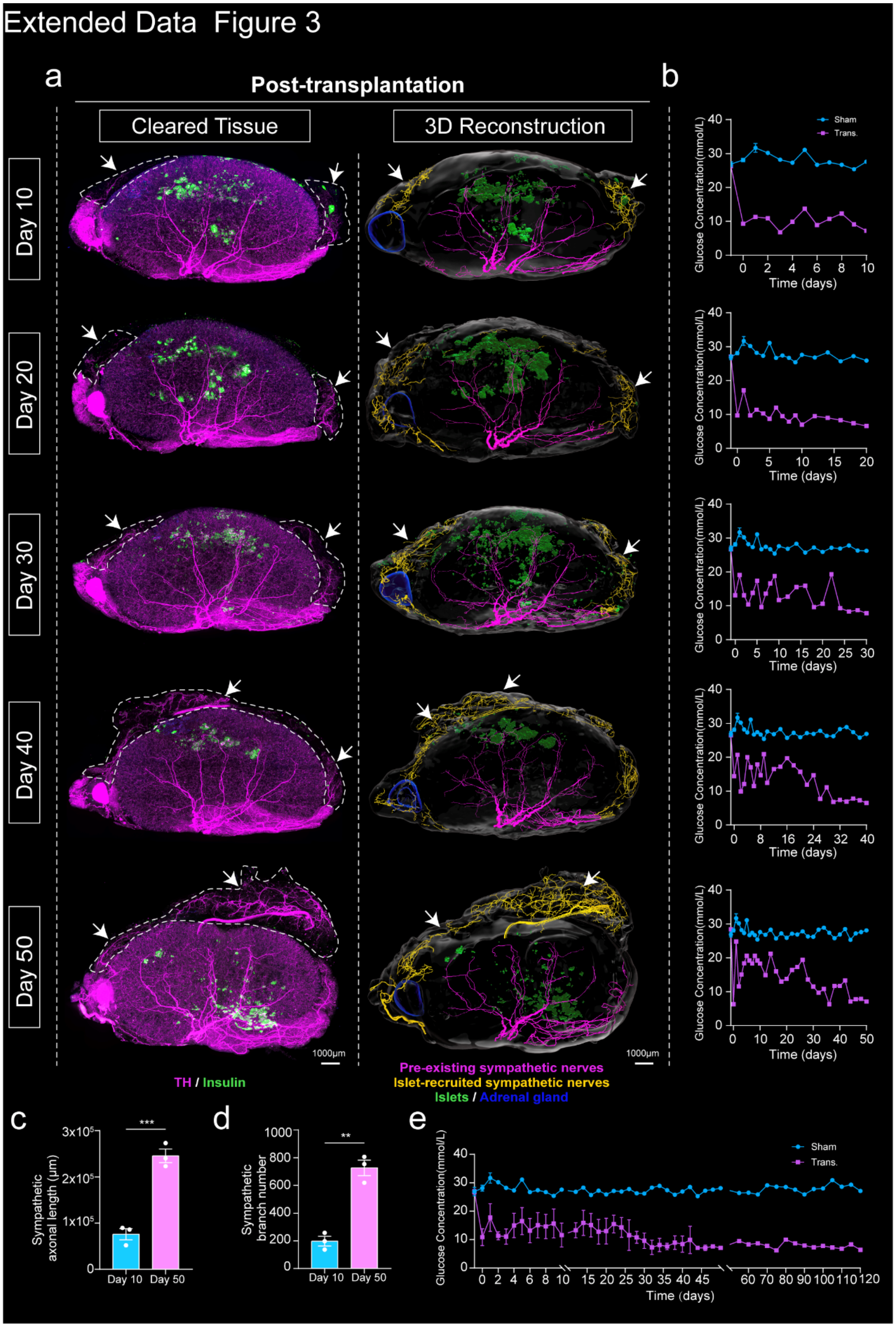
Extended Data Figure 3
a
Post-transplantation
Cleared Tissue
3D Reconstruction
Day 10
Day 20
Day 30
Day 40
Day 50
1000μm
TH / Insulin
Pre-existing sympathetic nerves
Islet-recruited sympathetic nerves
Islets / Adrenal gland
b
Sham
Trans.
Glucose Concentration(mmol/L)
Time (days)
c
Sympathetic axonal length (μm)
***
Day 10
Day 50
d
Sympathetic branch number
**
e
Sham
Trans.
Glucose Concentration(mmol/L)
Time (days)

**Extended Data Figure 3. Viability of transplanted pancreatic islets and their recruitment of sympathetic nerves.**

**a,** Representative images illustrating the progressive recruitment of sympathetic fibers toward transplanted islets at multiple time points (days 10, 20, 30, 40, and 50) post-transplantation. Left, whole-kidney tissue clearing images showing sympathetic nerves (TH, magenta) and transplanted islets (insulin, green); right, corresponding 3D reconstructions depicting pre-existing renal sympathetic nerves (magenta), islet-recruited sympathetic nerves (yellow), transplanted islets (green), and the adrenal gland (blue). White arrowheads indicate sympathetic nerves attracted toward islets. Scale bars: 1000 µm. Supplementary Videos 5-9.

**b,** Random blood glucose levels were measured at the indicated time points.

**c, d,** Quantification of sympathetic nerve fiber length (c) and branching number (d) at days 10 and 50 post-transplantation (n = 3 per group). Data are presented as mean ± SEM.

**e,** Summary of random blood glucose levels measured at each indicated time point.

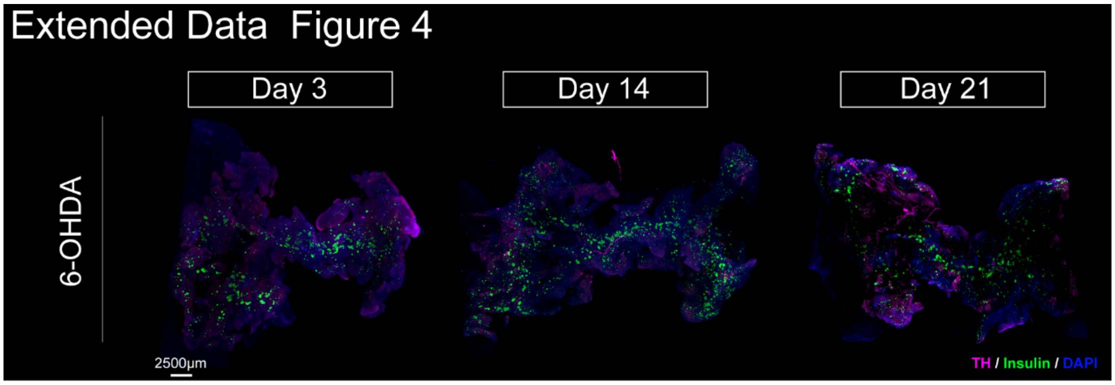


**Extended Data Figure 4. Validation of efficiency and specificity of sympathetic nerve ablation.**

Representative whole-pancreas images demonstrating successful sympathetic denervation at days 3, 14, and 21 after intrapancreatic injection of 6-OHDA, indicated by the near-complete absence of sympathetic fibers (TH, magenta). Pancreatic islets labeled by insulin (green), nuclei labeled by DAPI (blue). Scale bars: 2500 µm. See Supplementary Videos 12-14.

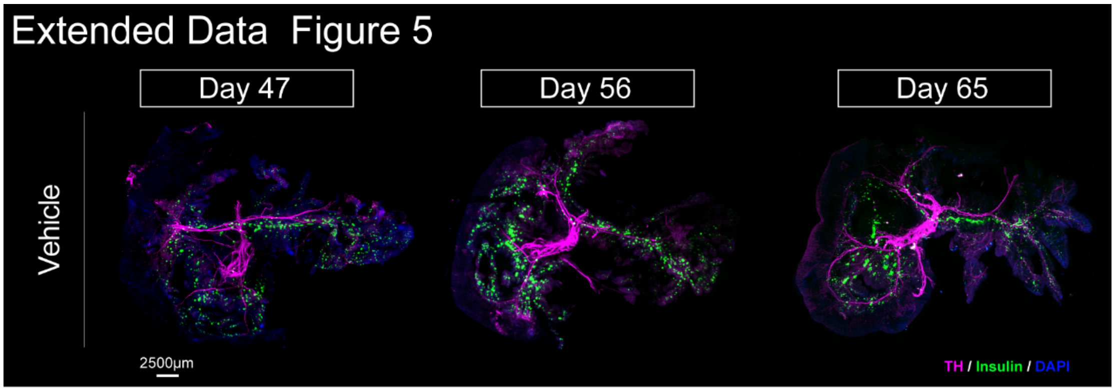


**Extended Data Figure 5. Effects of vehicle injection on sympathetic nerves.**

Control mice injected with vehicle solvents exhibited sympathetic innervation (TH, magenta). Pancreatic islets labeled by insulin (green), nuclei labeled by DAPI (blue). Scale bars: 2500 µm. See Supplementary Videos 21-23.

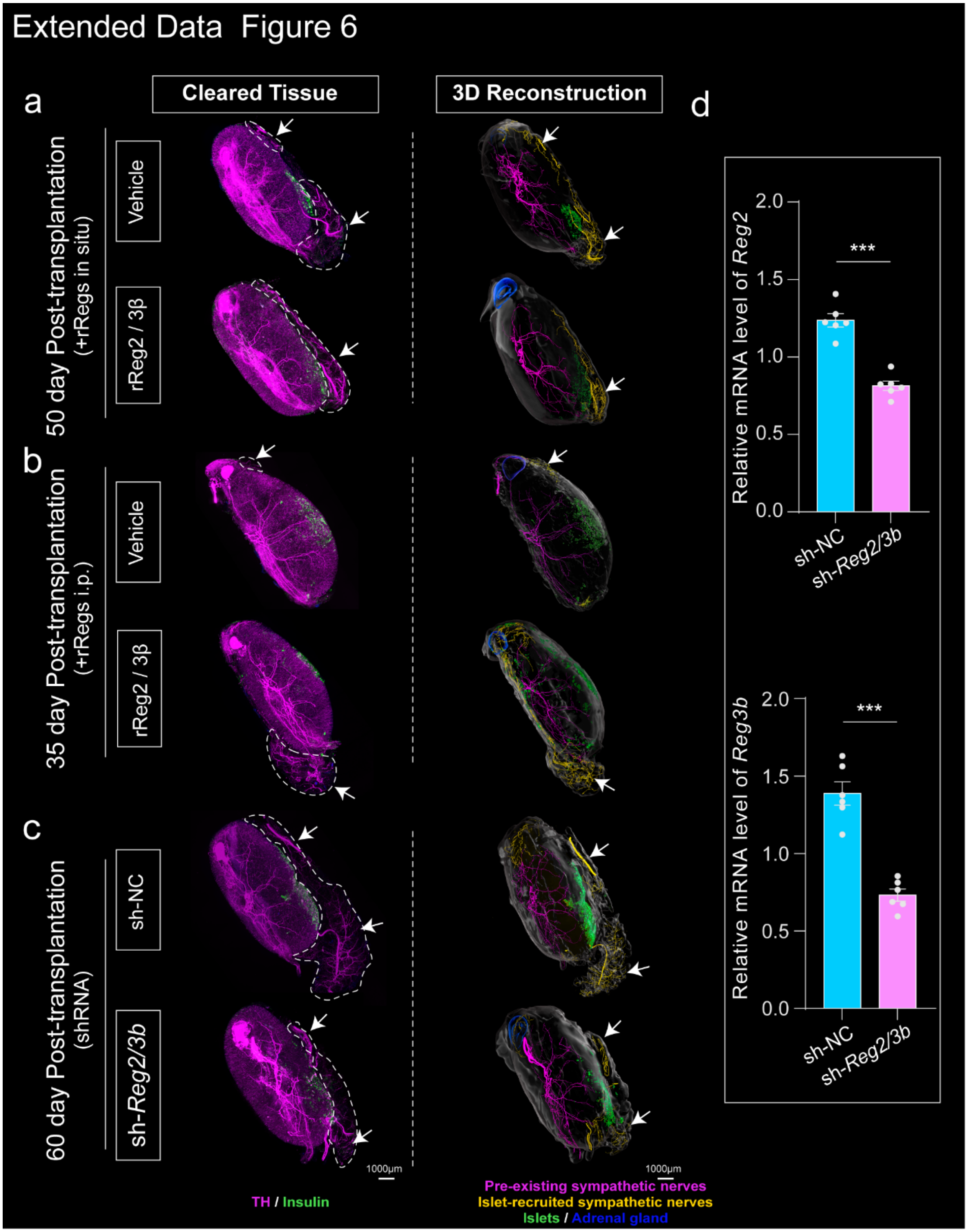
Extended Data Figure 6
a
b
c
d
Cleared Tissue
3D Reconstruction
50 day Post-transplantation
(+rRegs in situ)
35 day Post-transplantation
(+rRegs i.p.)
60 day Post-transplantation
(shRNA)
Vehicle
rReg2 / 3β
sh-NC
sh-Reg2/3b
1000μm
TH / Insulin
Pre-existing sympathetic nerves
Islet-recruited sympathetic nerves
Islets / Adrenal gland
Relative mRNA level of Reg2
Relative mRNA level of Reg3b
2.0
1.5
1.0
0.5
0.0
***
sh-NC
sh-Reg2/3b

**Extended Data Figure 6. Additional validation and 3D views of sympathetic innervation around transplanted islets.**

**a-c,** Representative 45° angled 3D images of whole-kidney cleared samples corresponding to experimental conditions in Figure 5a. Left: cleared kidney images showing sympathetic nerves (TH, magenta) and transplanted islets (insulin, green). Right: corresponding 3D reconstructions illustrating pre-existing renal sympathetic nerves (magenta), islet-recruited sympathetic nerves (yellow), transplanted islets (green), and adrenal gland (blue). White arrowheads indicate nerves attracted toward islets. Scale bars: 1000 µm.

**d,** Quantitative RT-qPCR analysis confirming efficient knockdown of *Reg2* and *Reg3b* in pancreatic islets following lentiviral-mediated shRNA transduction. Data represent mean ± SEM from n = 6 independent experiments; *$P < 0.05$, **$P < 0.01$, ***$P < 0.001$, two-tailed Student's t-test.

**Table S1.**

Antibody list

| Antigene | Source | Catalogue number | Antibody dilution |
|---|---|---|---|
| TH | abcam | AB112 | 1:500 |
| Insulin | R&D system | MAB1417 | 1:500 |
| CGRP | abcam | AB36001 | 1:500 |
| VACHT | Synaptic system | 139103 | 1:500 |
| CD31-A647 | Jarvisbio | AB152008 | 15-20ug/20g |
| DAPI | Invitrogen | D1306 | 1:500 |
| Donkey anti-Rabbit lgG (H+L) Highly Cross-Adsorbed Secondary Antibody, Alexa Fluor 647 | Invitrogen | A31573 | 1:500 |
| Donkey anti-Rat lgG (H+L) Highly Cross-Adsorbed Secondary Antibody, Alexa Fluor Plus 488 | Invitrogen | A48269 | 1:500 |
| Donkey anti-Goat lgG (H+L) Cross-Adsorbed Secondary Antibody, Alexa Fluor 555 | Invitrogen | A21432 | 1:500 |
| Donkey anti-Rabbit lgG (H+L) Highly Cross-Adsorbed Secondary Antibody, Alexa Fluor 555 | Invitrogen | A31572 | 1:500 |

**Table S2.**

Target Sequences of Lentivirus-Delivered shRNAs

| Gene | Gene ID | TargetSeq |
|---|---|---|
| NC | NA | CCTAAGGTTAAGTCGCCCTCG |
| Reg2 | NM_009043.2 | GCAGGTCACCTGGTGTCAATA |
| Reg2 | NM_009043.2 | GCCTATGGTTCCTACTGTTAT |
| Reg2 | NM_009043.2 | GCTCCCTATTTCTCTTCAAGT |
| Reg3b | NM_011036.1 | ATGACGTGATGAATTACTTTA |
| Reg3b | NM_011036.1 | GCCCTATGTCTGCAAATTTAC |
| Reg3b | NM_011036.1 | ATATACCCTCCGCACGCATTA |

**Table S3.**

Primers for RT-qPCR

| | |
|---|---|
| M-mReg2-Forward | ACGCCTATGGTTCCTACTGTTA |
| M-mReg2-Reverse | AGTGCCAACGACGGTTACT |

| | |
|---|---|
| M-mReg3b-Forward | GGCTTCATTCTTGTCCTCCAT |
| M-mReg3b-Reverse | AGCACGGTCTAAGGCAGTAG |
| M-Actb-Forward | GGCACCACACCTTCTACAATG |
| M-Actb-Reverse | CCTCGTAGATGGGCACAGTG |

**Supplementary Videos.**

**Supplementary Video 1**: 3D visualization of sympathetic nerve networks precisely innervating the pancreatic islets.

**Supplementary Video 2**: 3D visualization of the distribution of parasympathetic nerves and pancreatic islets in the pancreas.

**Supplementary Video 3**: 3D visualization of the distribution of sensory nerves and pancreatic islets in the pancreas.

**Supplementary Video 4:** Comprehensive 3D visualization of sympathetic nerves innervating transplanted islets and sham-operated control at day 120 post-transplantation.

**Supplementary Video 5:** 3D visualization of sympathetic nerve growth toward transplanted islets at day 10 post-transplantation.

**Supplementary Video 6:** 3D visualization of sympathetic nerve growth toward transplanted islets at day 20 post-transplantation.

**Supplementary Video 7:** 3D visualization of sympathetic nerve growth toward transplanted islets at day 30 post-transplantation.

**Supplementary Video 8:** 3D visualization of sympathetic nerve growth toward transplanted islets at day 40 post-transplantation.

**Supplementary Video 9:** 3D visualization of sympathetic nerve growth toward transplanted islets at day 50 post-transplantation.

**Supplementary Video 10:** Severely disrupted pancreatic sympathetic innervation at day 140 after STZ-induced chronic islet injury.

**Supplementary Video 11:** Pancreatic sympathetic innervation at day 140 after i.p. injection of vehicle (control for Supplementary Video 10).

**Supplementary Video 12:** Ablation of pancreatic sympathetic nerves at day 3 after intrapancreatic 6-OHDA injection.

**Supplementary Video 13:** Ablation of pancreatic sympathetic nerves at day 14 after intrapancreatic 6-OHDA injection.

**Supplementary Video 14:** Ablation of pancreatic sympathetic nerves at day 21 after intrapancreatic 6-OHDA injection.

**Supplementary Video 15:** Regeneration of pancreatic sympathetic nerves at day 47 post-ablation.

**Supplementary Video 16:** Regeneration of pancreatic sympathetic nerves at day 56 post-ablation.

**Supplementary Video 17:** Regeneration of pancreatic sympathetic nerves at day 65 post-ablation.

**Supplementary Video 18:** Lack of sympathetic nerve regeneration at day 47 post-ablation following islet injury.

**Supplementary Video 19:** Lack of sympathetic nerve regeneration at day 56 post-ablation following islet injury.

**Supplementary Video 20:** Lack of sympathetic nerve regeneration at day 65 post-ablation following islet injury.

**Supplementary Video 21:** Maintenance of sympathetic innervation in solvent-treated controls at day 47 post-injection.

**Supplementary Video 22:** Maintenance of sympathetic innervation in solvent-treated controls at day 56 post-injection.

**Supplementary Video 23:** Maintenance of sympathetic innervation in solvent-treated controls at day 65 post-injection.

**Supplementary Video 24:** Enhanced sympathetic nerve growth toward transplanted islets following local administration of rReg2 and rReg3β.

**Supplementary Video 25:** Enhanced sympathetic nerve regeneration following i.p. injection of rReg2 and rReg3β.

**Supplementary Video 26:** Impaired sympathetic nerve regeneration following lentiviral knockdown of *Reg2* and *Reg3b* in transplanted islets.

**Supplementary Video 27:** Sympathetic innervation of transplanted islets at day 50 following combined local and systemic administration of Reg2 and Reg3β.
